# Hippocampal area CA2 controls seizure dynamics, interictal EEG abnormalities and social comorbidity in mouse models of temporal lobe epilepsy

**DOI:** 10.1101/2023.01.15.524149

**Authors:** Christos Panagiotis Lisgaras, Azahara Oliva, Sam Mckenzie, John LaFrancois, Steven A. Siegelbaum, Helen E. Scharman

## Abstract

Temporal lobe epilepsy (TLE) is characterized by spontaneous recurrent seizures, abnormal activity between seizures, and impaired behavior. CA2 pyramidal neurons (PNs) are potentially important because inhibiting them with a chemogenetic approach reduces seizure frequency in a mouse model of TLE. However, whether seizures could be stopped by timing inhibition just as a seizure begins is unclear. Furthermore, whether inhibition would reduce the cortical and motor manifestations of seizures are not clear. Finally, whether interictal EEG abnormalities and TLE comorbidities would be improved are unknown. Therefore, real-time optogenetic silencing of CA2 PNs during seizures, interictal activity and behavior were studied in 2 mouse models of TLE. CA2 silencing significantly reduced seizure duration and time spent in convulsive behavior. Interictal spikes and high frequency oscillations were significantly reduced, and social behavior was improved. Therefore, brief focal silencing of CA2 PNs reduces seizures, their propagation, and convulsive manifestations, improves interictal EEG, and ameliorates social comorbidities.

**HIGHLIGHTS:**

- Real-time CA2 silencing at the onset of seizures reduces seizure duration
- When CA2 silencing reduces seizure activity in hippocampus it also reduces cortical seizure activity and convulsive manifestations of seizures
- Interictal spikes and high frequency oscillations are reduced by real-time CA2 silencing
- Real-time CA2 silencing of high frequency oscillations (>250Hz) rescues social memory deficits of chronic epileptic mice

## INTRODUCTION

Temporal lobe epilepsy (TLE) is the most common type of epilepsy in adults^1^ characterized by recurrent seizures and an increased risk of comorbidities such as memory deficits.^2^ Anti-seizure medications fail to control seizures in more than 30% of patients with epilepsy^3^ highlighting the need for a better understanding of underlying mechanisms so that therapeutic strategies can be improved. Moreover, epilepsy is associated with a number of behavioral and psychiatric comorbidities, further reducing the quality of life of affected individuals.^4^

The hippocampus is one of the primary brain areas affected in TLE and alterations in hippocampal function are a strong candidate for TLE cognitive comorbidities. Indeed, brain autopsies from many patients with TLE have shown extensive neuronal damage in dentate gyrus (DG) hilus, CA3 and CA1 and sparing of DG granule cells and CA2 pyramidal neurons (PNs).^5^ This resistance of the DG and CA2 neurons to neuronal loss in TLE has been proposed to make these areas critical to seizure propagation through the epileptic hippocampus.^6–8^ However, relatively few studies have directly tested the idea that area CA2 regulates seizures in TLE.

Our prior work in a mouse model of TLE has shown that CA2 PNs become hyperexcitable after pilocarpine-induced status epilepticus and that prolonged chemogenetic CA2 silencing for three-week periods reduces seizure frequency.^8^ However, as prolonged CA2 silencing can lead to secondary changes that might indirectly alter circuit function, and that may even paradoxically increase excitability of other hippocampal regions,^9^ the acute, direct role that CA2 plays in seizure activity remains uncertain. Moreover, it is unclear whether alterations in CA2 function may contribute to behavioral comorbidities associated with TLE.

This possibility that abnormal CA2 function in TLE may contribute to behavioral comorbidities is suggested by recent findings that CA2 normally plays a critical role in social memory formation, consolidation and recall.^10,11^ Hippocampal area CA2 synchronizes sharp wave ripples (SWRs),^12^ a form of population activity observed during slow wave sleep that is essential for the consolidation of social experience.^13^ Notably, disruption of CA2 SWRs leads to social memory deficits during social memory recall.^13^ As many patients with TLE show social comorbidities^14,15^ for which there currently is no available treatments, manipulations that target abnormal electrical activity in CA2 may provide a novel approach to treating social comorbidity.

Here we used *in vivo* electrophysiological recordings from the CA2 subfield and closed-loop optogenetics to inhibit CA2 selectively and acutely only during seizures. This approach allowed us to directly test the role of CA2 PNs in controlling seizure dynamics. We also asked whether CA2 PNs contribute to abnormal electrical activity between seizures, and we thus tested its role in the generation of interictal spikes (IIS) and high frequency oscillations (HFOs>250Hz). Last, we asked whether silencing abnormal electrical activity during the consolidation phase of social experience could improve social memory performance in chronic epileptic mice.

Our data provide novel insights into the direct role of CA2 activity in several major pathophysiological hallmarks of TLE and further suggest that area CA2 could become a new therapeutic target for treating TLE and associated social comorbidities.

## MATERIALS AND METHODS

### I. Animals

All experimental procedures were performed in accordance with the NIH guidelines and approved by the Institutional Animal Care and Use Committee (IACUC) at the Nathan Kline Institute. Male Amigo2-Cre^+/-^ mice (stock # 030215; The Jackson Laboratory) from the Siegelbaum laboratory (stock # 000664 C57BL/6J background; The Jackson Laboratory) were bred with wild-type (WT) female C57BL/6N mice (stock #B6-F/B6-M, Taconic) and backcrossed with the C57BL/6N line for several generations before use. For all experiments, we used adult male and female Amigo2-Cre^+/-^ mice that were 8-12 weeks-old at the start of the experiments. All mice were fed Purina 5001 (W.F. Fisher), with water *ad libitum*. Cages were filled with corn cob bedding and there was a 12 hr light:dark cycle (7:00 a.m. lights on, 7:00 p.m. lights off). Genotyping was performed by the Mouse Genotyping Core Laboratory at NYU Langone Medical Center.

### II. Epilepsy models

We used animals that developed chronic convulsive seizures after pilocarpine (PILO)- or intrahippocampal kainic acid (IHKA)-induced status epilepticus (SE). For PILO-SE induction,^8,16,17^ mice were injected subcutaneously (s.c.) with the muscarinic antagonist scopolamine methyl nitrate (1 mg/kg, s.c.; Sigma Aldrich) to reduce the peripheral effects of PILO. The β2 adrenergic agonist terbutaline hemisulfate (1 mg/kg, s.c.; Sigma Aldrich) was injected to support respiration. Ethosuximide (150 mg/kg, s.c.; Sigma Aldrich) was administered to reduce the occurrence of brainstem seizures, which often lead to mortality.^18^ SE was induced by injecting pilocarpine hydrochloride (250 mg/kg, s.c.; Sigma Aldrich), and after 2 hrs, mice were injected with diazepam (5 mg/kg, s.c.; Hospira) to reduce SE severity and 1 mL (s.c.) of lactated Ringer’s (Aspen Veterinary Resources) for hydration.

A separate cohort of mice was treated with IHKA to induce epilepsy as described previously.^19^ In brief, mice were anesthetized with 3% isoflurane (Henry Schein) and transferred to a stereotaxic apparatus (World Precision Instruments). One burr hole was drilled above the left hippocampus (−2 mm anterior-posterior [A-P] to Bregma, −1.25 mm medial-lateral [M-L]). A 0.5 mL Hamilton syringe (#7001, Hamilton) was lowered from the brain surface 1.6 mm into the left hippocampus (−1.6 mm dorsal-ventral [D-V]) and 100 nL of 20 mM KA was injected over 5 min. The needle remained in place for an additional 5 min to prevent backflow and the incision was closed with Vetbond (3M). Next, mice were monitored for SE defined as severe and continuous convulsive seizures^20^ lasting for several hours.

### III. Viral injections

PILO- and IHKA-SE mice were injected with virus 4 weeks after epilepsy induction when they already show spontaneous seizures.^8,17,19^ Mice were anesthetized with 3% isoflurane and then transferred to a stereotaxic apparatus where anesthesia was maintained with 1-2% isoflurane. Next, buprenorphine (0.05 mg/kg) was injected s.c. to reduce discomfort. One burr hole was drilled and 200 nL of adeno-associated virus carrying the inhibitory opsin Arch3.0 (AAV2/5-EF1a-DIO-eArch3.0-eYFP) or control flurophore (AAV2/5-EF1a-DIO-eYFP) were injected bilaterally to the following coordinates (−2.0 mm A-P, ±1.8 mm M-L, −1.7 mm D-V). The needle was then left in place for another 10 min and it was slowly retracted to avoid backflow of the injected virus.

### IV. Electrode implantation

Three weeks following viral injections mice were implanted with electrodes. Mice were anesthetized with 3% isoflurane and then maintained with 1-2% isoflurane. Next, buprenorphine (0.05 mg/kg, s.c.) was injected to reduce discomfort. Two burr holes were drilled above cerebellar region and subdural screw electrodes were placed and stabilized using dental cement (Lang Dental) to serve as a reference (−5.7 mm A-P, +1.25 mm M-L) and a ground (−5.7 mm A-P, −1.25 mm M-L). Next, a burr hole was drilled over the left hippocampus and one optrode (200 um optical fiber coupled to a 50 um wire electrode) was implanted unilaterally in the left dorsal CA2 (−1.94 mm A-P, −2.2 mm M-L, −1.8 mm D-V) and a screw electrode over right frontal cortex (−0.5 mm A-P, +1.5 mm M-L). Closed-loop optogenetic experiments were conducted at least 1 week after EEG surgery to allow for recovery from surgery.

### V. Closed-loop optogenetic experiments

For EEG recording, mice were housed in a 21 cm x 19 cm square transparent plexiglass cage with access to food and water. A pre-amplifier (16-channel headstage with accelerometer, Intan Techologies) was connected to the implant and then to a rotatory joint (Doric Lenses) via a lightweight cable to allow unrestricted movement of the mouse. In addition, a patch optical cable was connected to the implant through the same rotatory joint and then to a 532 nm laser source (Opto Engine). EEG signals were recorded at 2 kHz sampling rate using a bandpass filter (0.1-500 Hz) with an RHD interface board (Intan Techologies). High frame rate video (>30 frames/sec) was recorded simultaneously using an infrared camera (ac2000, Basler).

#### Online detection and interruption of seizures

For real-time seizure disruption, the EEG signal from the depth electrode in CA2 was routed to a real-time programmable processor (CED 1401, UK) through an analog port. The processor used a real-time sliding window of 2 sec to calculate the rate of interictal spike activity occurring at the onset of seizures and then trigger 500-ms-long light pulses when seizure activity crossed and remained above the threshold. Thresholds for seizure detection was set manually for each animal during a baseline period of ~4 days preceding the start of closed-loop tests when spontaneous seizures were recorded. In control interventions, seizures received no light stimulation or light stimulation was delivered to flurophore-expressing mice. Closed-loop tests continued for 4-5 days during which a subset of seizures was stimulated or not stimulated and that was chosen at random.

#### Real-time disruption of interictal spikes

Interictal spikes (IIS) were detected online using an amplitude threshold of 5 standard deviations above the mean of the baseline signal. For detecting IIS, we used the EEG signal from the depth electrode implanted in CA2 and fed it to the same programmable processor mentioned above. Thresholds for IIS detection were determined based on a baseline session where the threshold for IIS detection were manually adjusted for each animal. Once an IIS was detected, a 500 ms square-shaped light pulse was delivered through the optrode implanted in CA2. In control stimulations, 500 ms light pulses were delivered in flurophore expressing mice or laser power was off when IIS were detected.

#### Real-time disruption of high frequency oscillations

For HFO disruption, the EEG signal from CA2 was band pass filtered in the 250-500 Hz frequency range using a hardware filter and the root mean square of the signal was calculated using an analog circuit. The signal was then fed to a programmable processor and amplitude thresholds (2 times above the SD of mean baseline) were manually adjusted for each animal before the start of the experiments. When the threshold was crossed, a 100 ms square-shaped light pulse was delivered through the optrode implanted in CA2. In control stimulations, light pulses of the same duration were delivered in flurophore expressing mice.

### VI. Behavioral experiments

For all behavioral experiments, mice were handled daily and accommodated to the experimenter, room where behavioral experiments took place and cables for 3 consecutive days leading to the task.

#### Sociability

To test sociability of mice, mice were placed in an (0.5 m x 0.5 m) arena in the presence of a conspecific confined in a wire cup cage and an empty cup cage placed on the opposite corner of the arena. The subject mouse was left to explore the arena and wire cup cages for 5 min and it was then returned to its homecage. During the task we used 8–12-week-old male mice as conspecifics. The position of the animal was tracked using an overhead camera (ac2000, Basler). We measured the total time that the subject mouse spent exploring the two wire cup cages by analyzing video records. Social exploration was defined by the time that the subject mouse spent in active exploration of the wire cup cages such as times that the subject mouse was sniffing, touching, and interacting with the confined mouse. The behavioral preference score was calculated as the differential interaction between the time spent exploring the wire cup cage where the conspecific was confined (N) and empty wire cup cage (E) divided by the total exploration time.

#### Social recognition memory

During habituation sessions, mice were exposed to the empty arena for 5 min followed by 5 min to the empty cup cages. Then mice were placed back to their homecage and after 1 hr they were exposed again to the arena containing the empty cup cages. Habituation sessions were done for 3 consecutive days leading to the task.

In the test day, mice were exposed for 5 min to the two novel stimulus animals in learning trial 1, 5 min to the same stimulus mice but in reversed position in learning trial 2 followed by 1 hr in home cage (during which HFOs were disrupted in the closed-loop cohort). After the 1 hr period in the homecage, mice were tested again in the recall test trial when one of the two stimulus mice (chosen at random) was presented along with a novel mouse. In the closed-loop cohort, mice were recorded with video-EEG for 3 days preceding the task and throughout the task to ensure they didn’t experience any seizures which would confound the behavioral test. Indeed, all 3 mice in the closed-loop cohort had no seizures during the 3-day recording period and no seizures occurred during the task. For each trial we measured the total time the subject mouse spent exploring the two wire cup cages where the stimulus mice were confined (S1, S2) in the two learning trials and differential time spent exploring the novel mouse (N) vs stimulus mice (S1 or S2) in the recall trail. The differential interaction was calculated using a discrimination index (DI) where DI= (t_N_ – t_S1_)/(t_N_ + t_S1_).

### VII. Quantification

#### Seizures

For offline seizure analyses, seizures were defined as sudden and rhythmic (>5 Hz) deflections in all EEG channels that lasted >10 sec and were at least 2 standard deviations (SD) above the baseline mean.^16,19^ The baseline mean was calculated from the 30 s prior to the event that was being considered to be a possible seizure. The threshold, 2 times the SD of the baseline mean, was chosen because it was adequate to differentiate seizures from normal EEG. Ten seconds was chosen because seizures in TLE typically last at least 10 sec and often are 20–60 sec^21^ As seizure onset we defined the time when the baseline of the left hippocampal lead exceeded 2 times the SD of the baseline mean. The end of a seizure (seizure termination) was defined as the time when high amplitude activity declined to less than 2 times the SD of the baseline mean. Seizure duration was calculated by subtracting the time of seizure termination from the time of seizure onset. Seizures were considered convulsive if the video record showed behaviors consistent with stages 3-5 on the Racine scale (stage 3, unilateral forelimb clonus; stage 4, bilateral forelimb clonus with rearing; stage 5, stage 4 followed by loss of posture.^20^ Seizures were considered non-convulsive when no stage 3-5 behaviors were detected in the video record, but the criteria for the EEG manifestation of the seizure were met. The time spent in convulsion was calculated based on video records and 3-axis acceleration during the EEG manifestation of a seizure. For spectral analyses we used the RippleLab application in MATLAB^22^ and time-frequency function. For power analyses we used Spike2 software.

#### Interictal spikes

IIS were defined using the same threshold as mentioned above for online IIS detection (5 standard deviations above the mean of the baseline signal). For amplitude analyses, we averaged all detected IIS from all mice used in the study. Averages were triggered from the time of online IIS detection and IIS amplitude is expressed as mean ± standard error of the mean (SEM).

#### High frequency oscillations

HFOs were defined as oscillations >250 Hz consistent with past studies in humans^23^ and animals.^16,19,24^ For offline HFO analyses we used the RippleLab application written in MATLAB ^22^ and the algorithm developed by Staba and colleagues.^25^ The amplitude of HFOs was calculated based on RMS amplitude of the 250-500 Hz filtered EEG signal when the laser power was on vs when the laser power was off. HFO duration was calculated using the same criteria we described previously.^16^ HFO power in the 250-500 Hz frequency range was calculated using Spike2 software.

### VIII. Statistics

Most data are presented as a mean ± SEM, unless otherwise stated. In box-and-whiskers plots the middle lines indicate medians; the bottom and top line of the box indicate the 25^th^ and 75^th^ quartiles, respectively and whiskers extend to the 5^th^ and 95^th^ percentile, respectively. Dots indicate individual data points. Statistical significance was set at p<0.05 and is denoted by asterisks on all graphs. Statistical comparisons that did not reach significance are designated by a “ns” (non-significant) symbol.

All statistical analyses were performed using Prism (Graphpad). To determine if data fit a normal distribution, the Shapiro-Wilk test was used. Comparisons of parametric data of two groups were conducted using unpaired or paired two-tailed Student’s t-tests. When data did not fit a normal distribution, non-parametric statistics were used. The non-parametric test to compare two groups was the Mann-Whitney U-test. For comparisons of more than 2 groups, one-way ANOVA was used when data were parametric and Kruskal-Wallis for non-parametric data. When a statistically significant main effect was found by ANOVA, Bonferroni post-hoc tests were used with corrections for multiple comparisons and for Kruskal-Wallis, Dunn’s post-hoc tests were used. For correlation analyses we used linear regression and the Pearson or Spearman correlation coefficient (r) in Graphpad.

## RESULTS

### Real-time CA2 silencing reduced hippocampal seizure duration and spectral power

To test whether acute CA2 PN silencing could disrupt spontaneous convulsive seizures, we induced chronic epilepsy in Amigo2-Cre mice, which express Cre-recombinase relatively selectively in CA2 PNs,^10^ using either pilocarpine (PILO)-^8,16,17^ or intrahippocampal kainic acid (IHKA)-induced^19^ SE. The mice were injected in CA2 with Cre-dependent AAV to express the inhibitory opsin Arch3.0 selectively in CA2 pyramidal neurons. We configured a closed-loop optogenetic protocol to detect spontaneous seizures in real-time and trigger light pulses starting at the onset of a subset of spontaneous seizures to inhibit CA2. Which seizures received light pulses, and which did not was decided in a random fashion. Light stimulation consisted of a series of 500 ms light pulses delivered to the left dorsal CA2 (Fig. 1A). In control interventions, seizures received no light pulses or light pulses were delivered in fluorophore-expressing mice.

**Figure 1:**
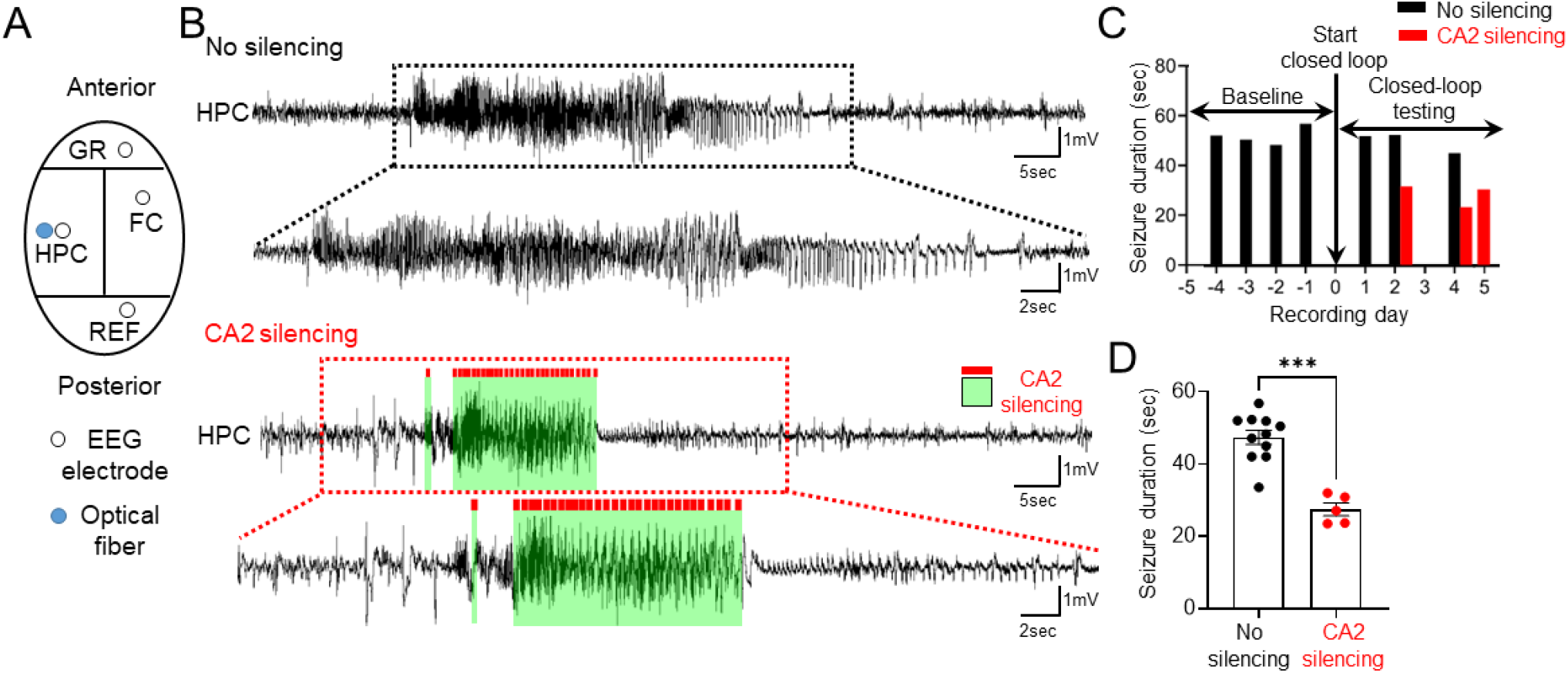
Real-time optogenetic silencing of CA2 PNs during seizures reduces hippocampal seizure duration. (A) EEG-recording diagram. Mice were implanted with an optrode in left dorsal CA2 and a screw electrode over right frontal cortex. HPC = hippocampus, FC = frontal cortex, GR = ground, REF = reference. (B) Example spontaneous seizures in an epileptic IHKA-SE mouse where CA2 PNs were not silenced (top) or silenced (bottom). The period of time that CA2 was silenced is denoted by green shading. For the top and bottom examples, dotted boxes surrounding some of the EEG record are expanded below. (C) Seizure duration before (baseline) and after the start of closed-loop tests. Each bar denotes the duration of a spontaneous seizure either during baseline or after closed loop testing began. Note when closed loop testing began, some seizures received no stimulation and had durations like seizures during the baseline (black). Other seizures received stimulation and had short durations (red). (D) Hippocampal seizure duration was significantly reduced upon CA2 silencing (both PILO and IHKA models) vs no silencing (paired Student’s t-test, t=10.6, df=4, p=0.0004).

Fig. 1A-B shows the electrode arrangement and a representative example of a seizure in an epileptic IHKA-SE mouse when CA2 PNs were not silenced (top) or were silenced (bottom). Example seizure durations before and after the start of closed-loop tests are shown in Fig. 1C. We found that seizure duration was significantly reduced upon CA2 PN silencing compared to when CA2 was not silenced (paired Student’s t-test, t=10.6, df=4, p=0.0004; Fig. 1D). In addition to the robust reduction in seizure duration, total EEG spectral power during hippocampal seizures was significantly reduced upon CA2 silencing (Fig. 2B) vs no silencing (Fig. 2A) in both TLE models (Fig. 2C, D).

**Figure 2:**
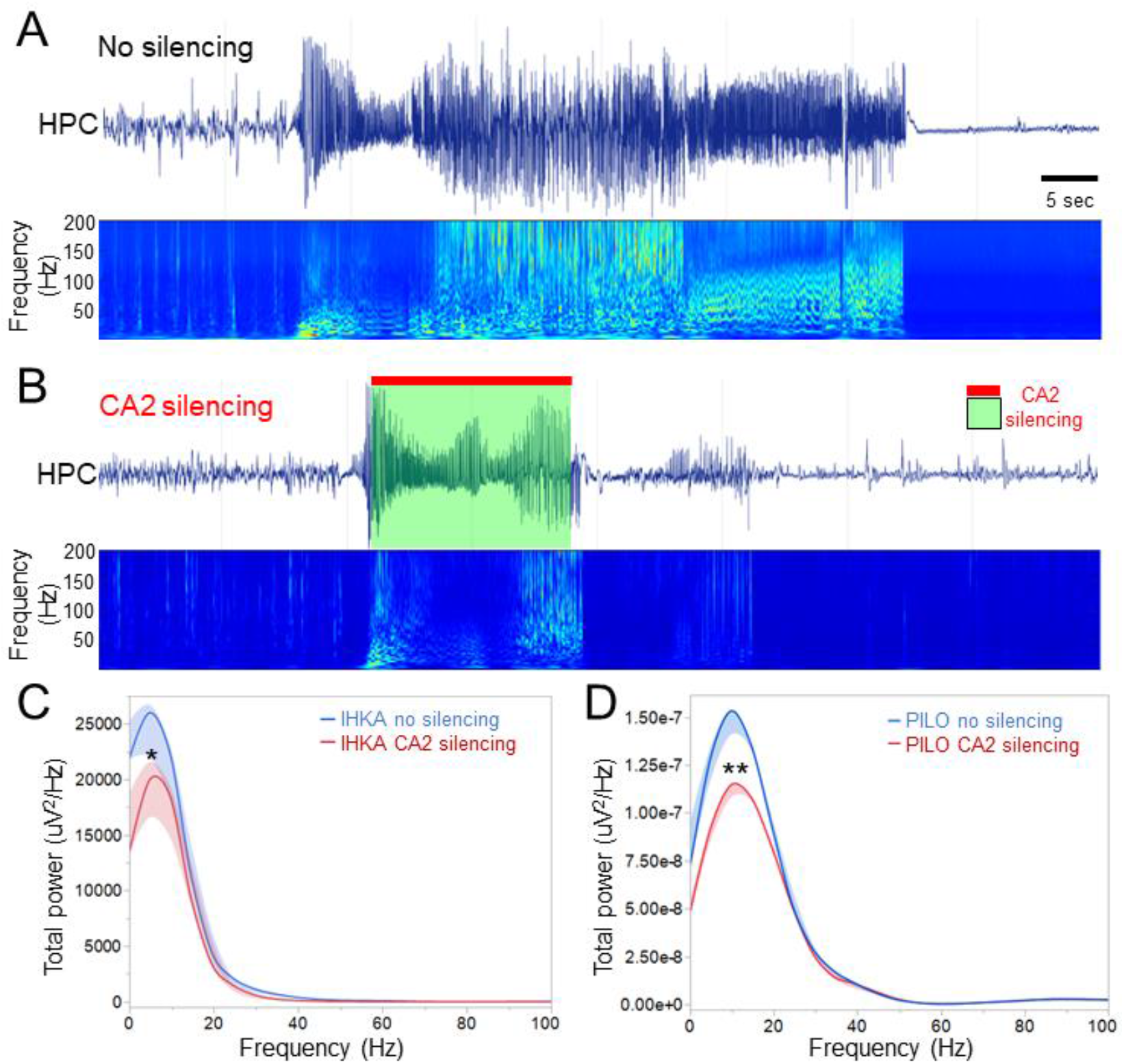
Seizure power in hippocampus is significantly reduced during CA2 silencing. (A) An example spontaneous seizure in an epileptic PILO-SE mouse where CA2 PNs were not silenced. Note robust spectral power in the 20-200Hz frequency range during the seizure. (B) Same as in A but for a seizure where CA2 PNs were silenced. Note reduced spectral power compared to seizure shown in A. (C) Spectral power was significantly reduced during CA2 silencing in IHKA-SE mice (p<0.05), n=3 IHKA mice. Blue = no silencing, Red = CA2 silencing. (D) Same as in C but for PILO-SE mice. Note that power was also significantly reduced upon CA2 silencing (p<0.01), n=3 PILO mice.

### CA2 silencing reduced cortical seizure power

An important characteristic of the CA2 circuitry is that area CA2 participates in a powerful excitatory cortico-hippocampal loop.^26,27^ This CA2 property is important as it would endow CA2 with the ability to potentially influence seizure propagation. To test whether CA2 silencing had a global rather than local inhibitory effect on seizures, we analyzed seizures recorded in the contralateral hemisphere from where CA2 PNs were silenced i.e., right frontal cortex (Fig. 3A, B). We found that cortical EEG spectral power during seizures was significantly reduced during CA2 silencing (p<0.01; Fig. 3D).

**Figure 3:**
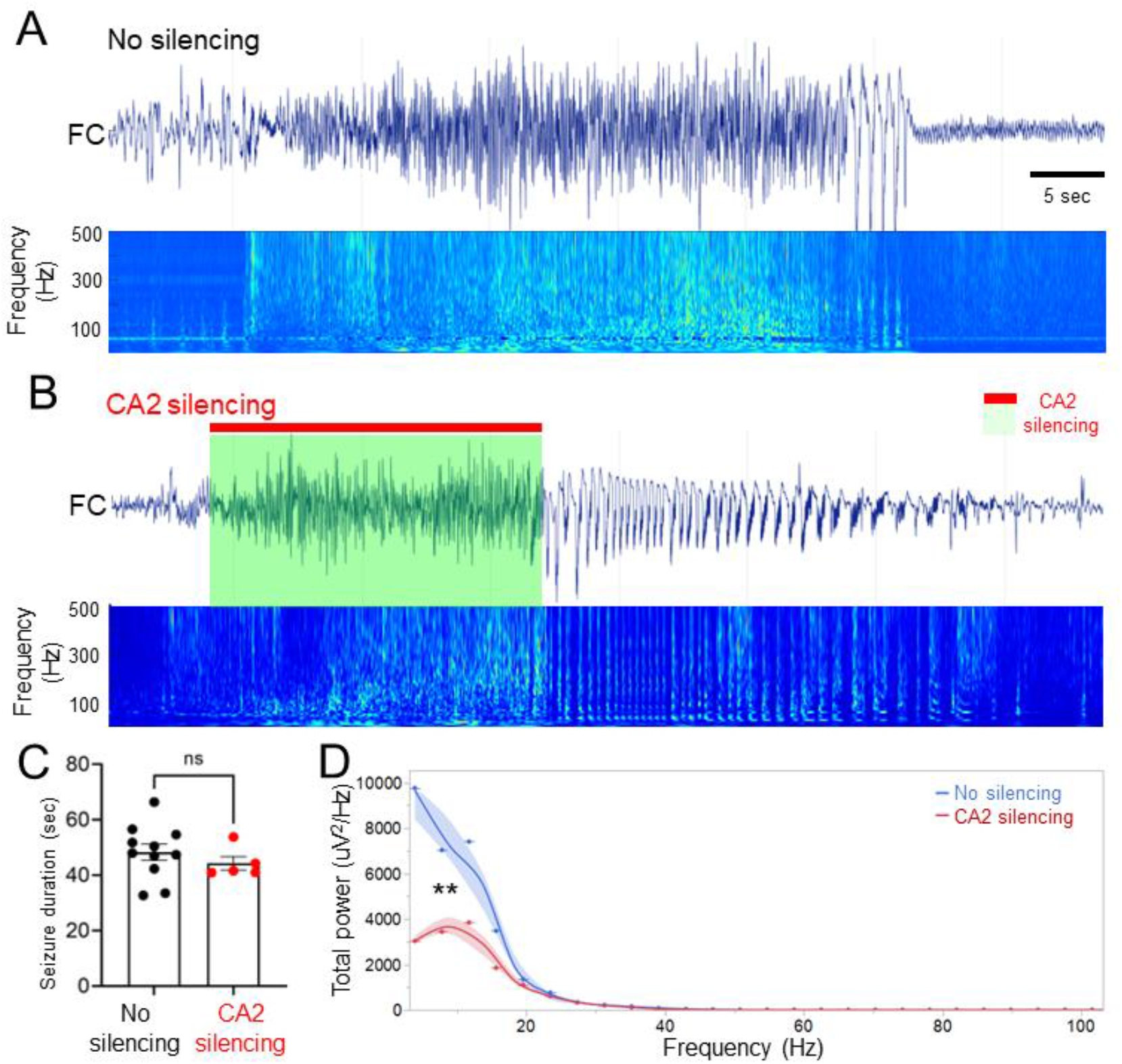
Closed-loop silencing of CA2 PNs reduces cortical EEG spectral power. (A) An example spontaneous seizure recorded from the right frontal cortex of a PILO-SE mouse when CA2 was not silenced. (B) Same as in A but for a seizure where CA2 PNs were silenced. (C) Cortical seizure duration was not significantly reduced upon CA2 PN silencing (both TLE models) vs no silencing (Wilcoxon signed rank test, p=0.31), n=5 mice (n=3 IHKA, n=2 PILO). (D) Cortical seizure power was significantly reduced when CA2 PNs were silenced vs no silenced (p<0.01).

However, cortical seizure duration was not significantly reduced during CA2 PN silencing (Wilcoxon signed rank test, p=0.31; Fig. 3C), indicating that CA2 inhibition does not influence the duration of remote electrographic seizure activity once the seizure generalized. Rather, CA2 PN silencing affects cortical spectral power of cortical seizure activity indicating a possible contribution of CA2 activity to the intensity of seizures.

### Real-time silencing of CA2 PNs during spontaneous seizures reduced convulsive behavior

The robust reduction in cortical seizure power during CA2 silencing prompted us to test whether CA2 silencing would also influence the behavioral severity of convulsive seizures. In both TLE models, spontaneous convulsive seizures were accompanied by robust convulsive behavior as measured by a 3-axis accelerometer (Fig. 4A) and verified by simultaneous video records. CA2 silencing during convulsive seizures significantly reduced the total time spent in convulsive behavior during the EEG manifestation of the seizure (paired Student’s t-test, t=8.4, df=6, p=0.001; Fig. 4B, C). In addition, whereas we found a positive significant correlation between hippocampal seizure duration and duration of convulsion when CA2 was spontaneously active (i.e., without light stimulation), (Pearson *r*=0.61, p=0.03; Fig. 4D), hippocampal seizure duration was no longer significantly correlated with the duration of convulsion in seizures when CA2 was silenced (Spearman *r*=-0.39, p=0.46; Fig. 4E). This suggests that without the participation of CA2 longer convulsive seizures were no longer accompanied by longer convulsions.

**Figure 4:**
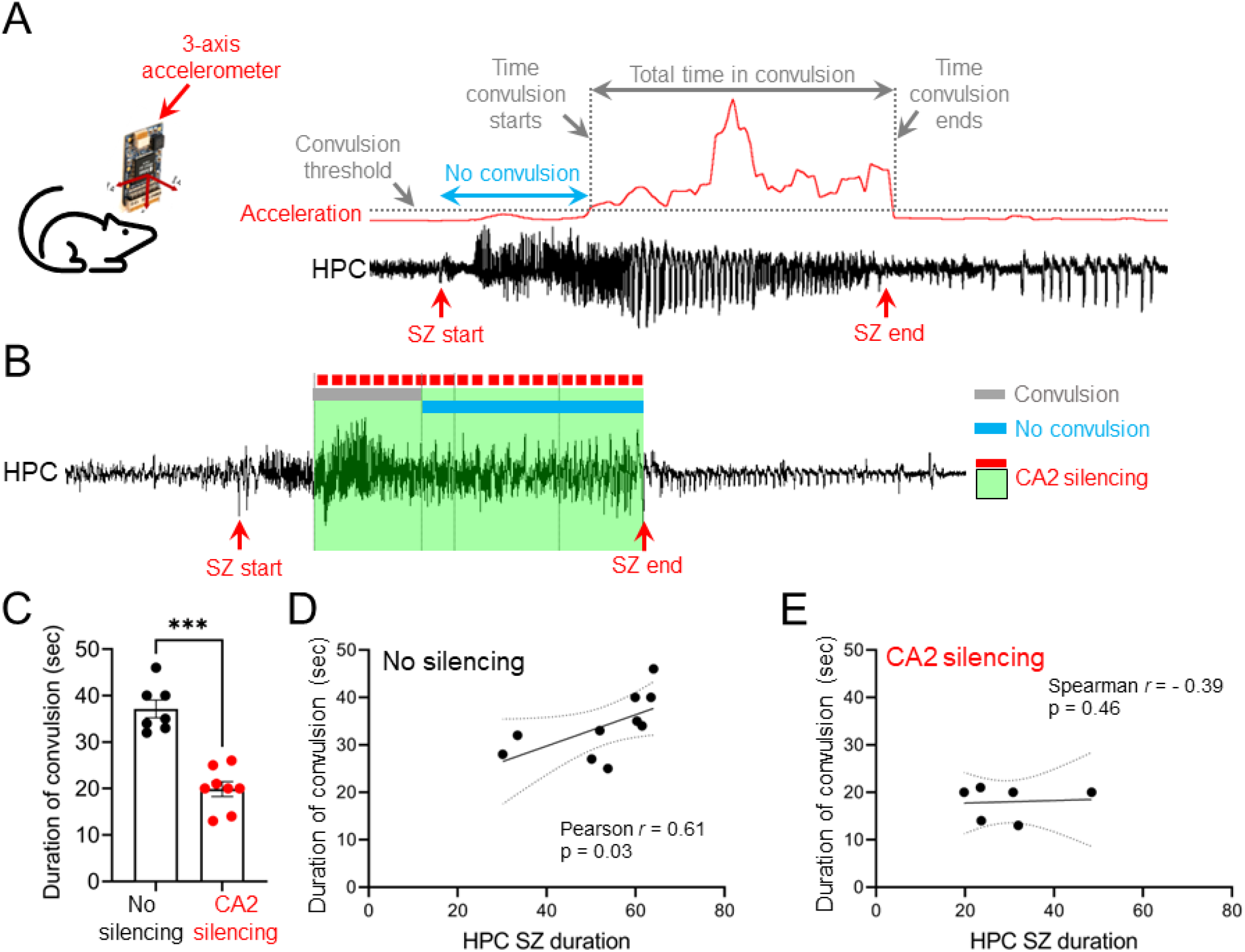
Time spent in convulsion is shortened during CA2 silencing. (A) An example of seizure and 3-axis acceleration measurements. Acceleration was measured in each of the three dimensions (x, y, z) and a composite acceleration was calculated and is shown by a red line. The time spent in convulsion was calculated based on a threshold as described in Methods. (B) An example seizure where CA2 PNs were silenced. The time spent in convulsion is shown with a grey bar and the time spent without convulsion with a blue line. The time that CA2 PNs were silenced is shown with green shading. Note that the majority of the EEG recorded seizure was accompanied by no convulsion. (C) The time spent in convulsion was significantly reduced when CA2 PNs were silenced vs no silencing (paired Student’s t-test, t=8.4, df=6, p=0.001), n=5 mice (n=3 IHKA, n=2 PILO). (D) Correlation between the duration of the convulsion and hippocampal seizure duration for seizures where CA2 PNs were not silenced. Note that there was a statistically significant positive correlation between the duration of the convulsion and hippocampal seizure duration (Pearson *r*=0.61, p=0.03), n=10 seizures from n=4 mice. (E) Same as in D but for seizures where CA2 PNs were silenced. Note the lack of a statistically significant correlation between the duration of the convulsion and hippocampal seizure duration (Spearman *r*=-0.39, p=0.46), n=6 seizures from n=3 mice.

### Interictal spikes are reduced in amplitude when CA2 PNs are silenced

Another important EEG abnormality in TLE is the presence of interictal spikes, brief large amplitude events caused by synchronous activation of a large population of neurons.^28^ IIS contribute to memory deficits in TLE^29^ and it has been suggested that they may originate in hippocampal area CA2.^30–32^ However, direct *in vivo* evidence for the emergence of IIS in CA2 is missing. To determine whether CA2 PNs contribute to IIS we detected IIS in real-time with the recording electrode in CA2 (Fig. 5A), and silenced CA2 PNs at the recording site as soon as IIS were detected there (Fig. 5B). We found that CA2 silencing truncated the spike component of the IIS at the point where the threshold was set for IIS detection (Fig. 5B). Indeed, IIS were significantly reduced in amplitude (p<0.001; Fig. 5C) and control optical stimulations (Fig. 5D, E) did not significantly reduce IIS amplitude (p>0.05; Fig. 5F), suggesting that the activity of CA2 PNs directly contributes to IIS.

**Figure 5:**
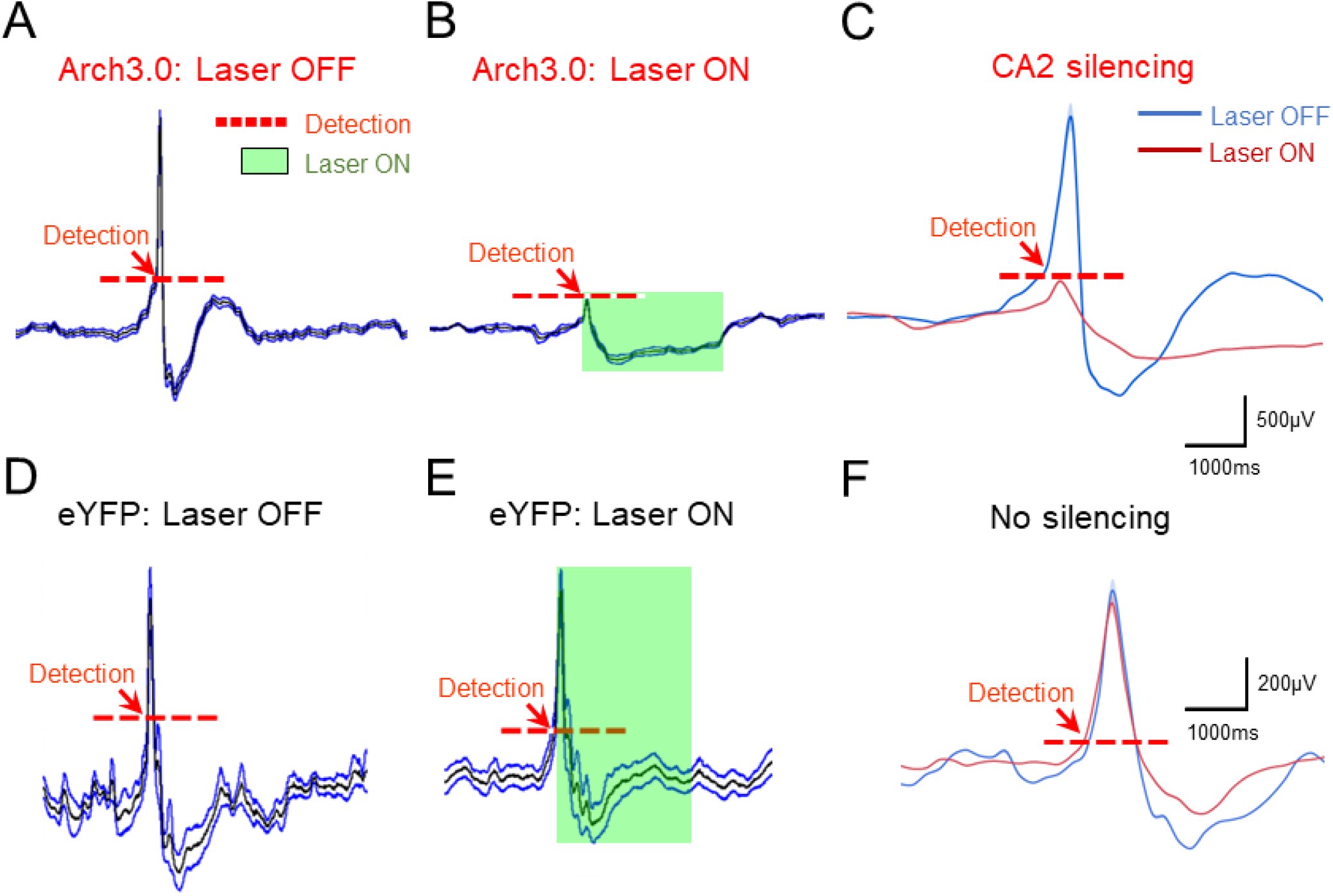
CA2 silencing at the time of IIS reduces IIS amplitude. (A) IIS average in Arch3.0-expressing mice when the laser power was off (n= 2 PILO, n=2 IHKA). Mean IIS amplitude is shown in black and SEM in blue. The time of IIS detection is denoted by a red arrow and the threshold used for IIS detection by a red dotted line. (B) Same as in A but the laser power was on (in the same 4 mice). Note robust reduction in IIS amplitude upon light stimulation. The time that the laser power was on is denoted by a green box. (C) Superimposition of traces shown in A and B. There was a statistically significant reduction in IIS amplitude when the laser power was on vs off (p<0.001). (D) IIS average in flurophore (eYFP)-expressing mice when the laser power was off (n= 2 PILO, n=1 IHKA). (E) Same as in D but the laser power was on (in the same 3 mice). (F) Superimposition of traces shown in D and E. Note that there were no statistically significant differences (p>0.05) in IIS amplitude during the period of time that laser power was on vs the same period of time that the laser power was off.

### CA2 inhibition reduces hippocampal high frequency oscillations

CA2 PNs are known to synchronize population activity during SWRs,^12^ but their contribution to abnormal HFOs such as those above 250 Hz remain unknown. To test whether CA2 PN activity contributes to HFOs we configured a closed-loop optogenetic protocol to detect HFOs in real-time and trigger brief (10 ms) light pulses at the time of HFO detection (Fig. 6A).

**Figure 6:**
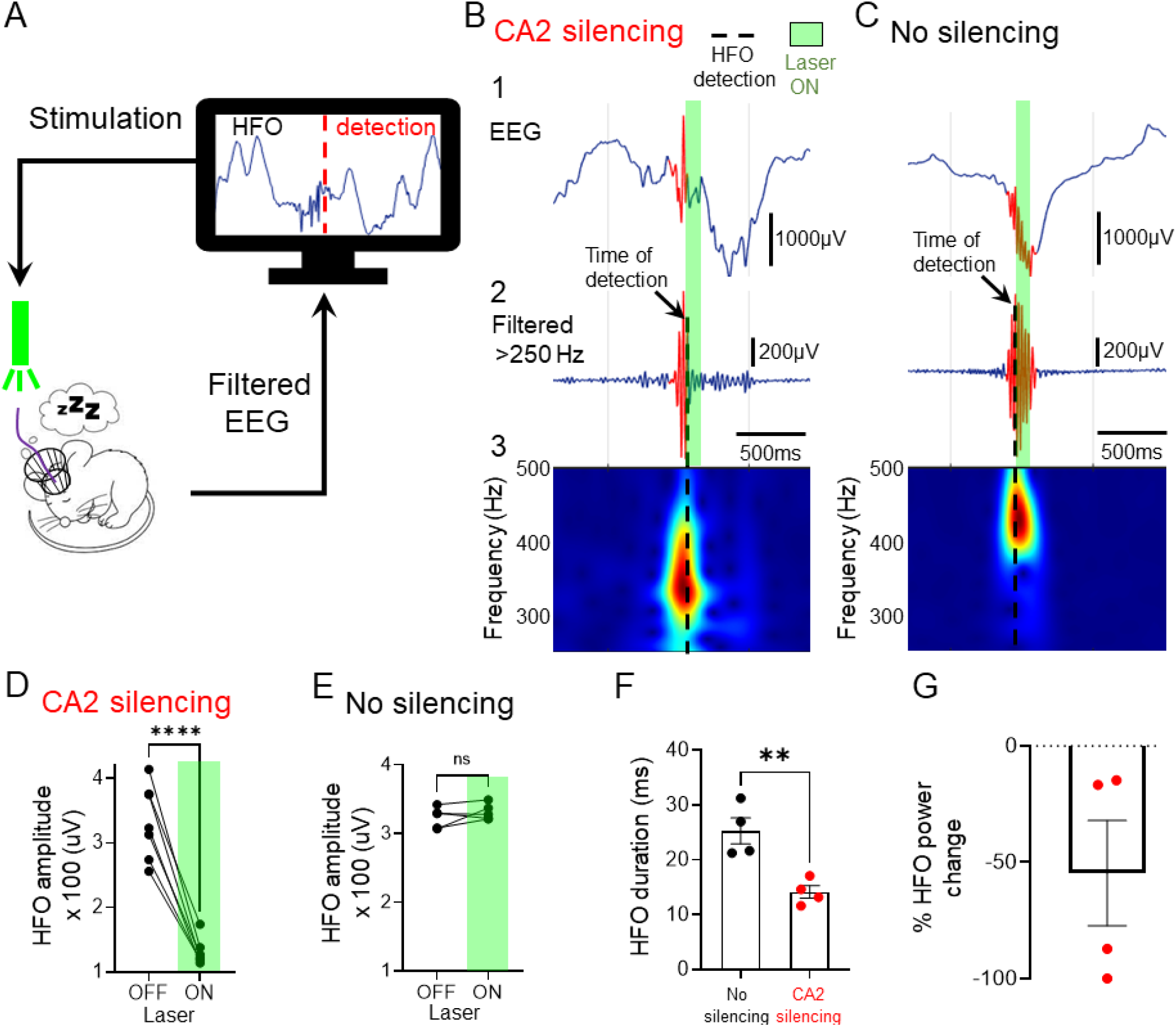
CA2 silencing at the time of HFOs reduces HFO amplitude, duration and power. (A) Schematic of the closed-loop system used for online disruption of HFOs during sleep. (B) An example HFO recorded in Arch3.0-expressing mouse. The time of HFO detection is denoted by a black arrow and the time that the laser power was on by green shading. Note reduced HFO power during light stimulation. Panel 1 shows raw EEG. Panel 2 shows the signal after filtering in the 250-500Hz which defines HFOs. Panel 3 is the spectrogram in the 250-500Hz frequency range. (C) Same as in B but for a flurophore expressing mouse. Note that HFOs power remains robust during light stimulation. (D) HFO amplitude for HFOs that received light stimulation (Laser ON) and those that did not (Laser OFF) in Arch3.0-expressing mice. HFO amplitude was significantly different between laser off and on (paired Student’s t-test, t=10.80, df=6, p<0.0001). (E) Same as in C but for HFOs in flurophore-expressing mice. HFO amplitude was not significantly different between laser off and on (paired Student’s t-test, t=1.26, df=4, p=0.27). (F) HFO duration is shown for HFOs that CA2 was not silenced or silenced. There was a statistically significant reduction in HFO duration when CA2 was silenced vs not silenced (paired Student’s t-test, t=6.04, df=6, p=0.009). (G)Percent change in HFO power upon CA2 silencing is shown (as compared to non-stimulated conditions). Interestingly, the two mice with the most reduced HFO power (over 75%) were IHKA mice and the two mice with less of a reduction in HFO power (<50%) were PILO mice.

CA2 silencing at the time of HFO detection aborted the HFO from continuing to occur (Fig. 6B) while had no effect in control stimulations (Fig. 6C). Indeed, HFO amplitude was significantly reduced by CA2 silencing in Arch3.0-expressing mice (paired Student’s t-test, t=10.80, df=6, p<0.0001; Fig. 6D) but not in eYFP flurophore-expressing mice (paired Student’s t-test, t=1.26, df=4, p=0.27; Fig. 6E). In addition to the robust reduction in HFO amplitude, HFO duration (paired Student’s t-test, t=6.04, df=6, p=0.009; Fig. 6F) was significantly reduced along with a marked reduction in HFO power compared to non-stimulated conditions (−54.7±22.6%; Fig. 6G).

### Impaired social memory in TLE is rescued by silencing CA2 during high frequency oscillations

As CA2 PN activity is essential for the encoding, consolidation and recall of social experience,^10,13^ we asked whether abnormal CA2 activity in the epileptic mice may lead to abnormal social behavior. This question is of clinical relevance as many patients with epilepsy show impaired social cognition.^14,15^ We first examined whether the epileptic mice have altered sociability, the normal preference of a mouse to explore another mouse compared to an empty chamber. We tested both normal (non-epileptic control) and epileptic mice in a two-choice paradigm where mice were placed in an arena containing two wire cup cages, one empty and the other containing a novel conspecific (Fig. 7A). We found that both normal and epileptic mice showed a significant preference for exploring the cage with the conspecific compared to the empty cage as suggested by significantly longer times spent around the conspecific vs the empty cage (Normal: paired Student’s t-test, t=6.2, df=3, p=0.008; Epileptic: paired Student’s t-test, t=8.4, df=3, p=0.003; Fig. 7B). When we quantified the behavioral preference score, we found no significant difference between normal and epileptic mice (unpaired Student’s t-test, t=1.04, df=4, p=0.35, n=3 mice per group; Fig. 7C). Importantly, total exploration times were not significantly different between normal and epileptic mice (unpaired Student’s t-test, t=1.3, df=6, p=0.24; Fig. 7D).

**Figure 7:**
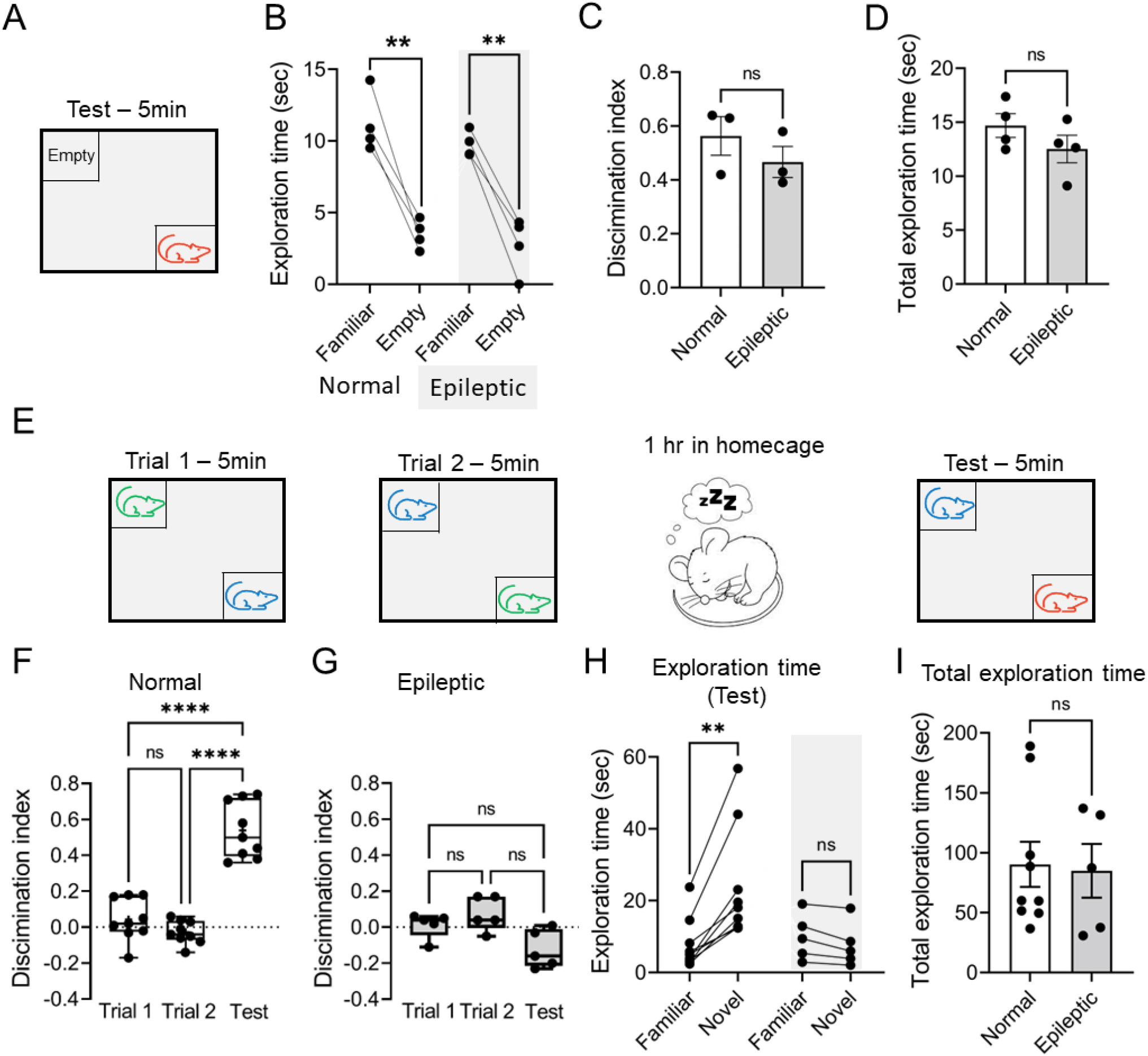
Social memory is impaired in chronic epileptic mice, but sociability is not. (A) Diagram of the sociability task. The subject mouse is presented with a familiar mouse and an empty cage and allowed to explore for 5 min. (B) Normal and epileptic mice showed a clear preference for the conspecific vs empty cage as showed by significant longer times spent exploring the conspecific vs empty cage (Normal: paired Student’s t-test, t=6.2, df=3, p=0.008; Epileptic: paired Student’s t-test, t=8.4, df=3, p=0.003). (C) The behavioral preference score was not significantly different between normal and epileptic mice (unpaired Student’s t-test, t=1.04, df=4, p=0.35, n=3 mice per group). (D) The total exploration time during the sociability task was not significantly different between normal and epileptic mice (unpaired Student’s t-test, t=1.3, df=6, p=0.24). (E) Diagram of the social discrimination task. The subject mouse is presented with 2 mice placed in the opposite corners of an arena for 5 min during trail 1. During trial 2 mice are switched in position and the subject mouse is kept in the arena for 5 min. Then the subject mouse is placed back to its homecage for 1 hr. After 1 hr the subject mouse is placed back in the arena where one mouse (familiar) is kept in its original position from trial 2 and a novel mouse is introduced to the opposite corner of the arena. (F) Non-epileptic controls (n=9) successfully recognize a novel from a familiar mouse at the recall test phase (One way ANOVA, F(5, 30)=32.17, p<0.0001). The middle lines on the box plot indicate medians; the bottom and top line of the box indicate the 25^th^ and 75^th^ quartiles, respectively and whiskers extend to the 5^th^ and 95^th^ percentile, respectively. (G) Epileptic mice (n=5) show impaired social recognition memory (One way ANOVA, F(2, 12)=2.28, p=0.14). (H) Normal mice showed a clear preference for the novel mouse compared to the familiar mouse, as showed by significantly longer times spent exploring the novel mouse (Wilcoxon signed rank test, p=0.003). Epileptic mice appeared to be impaired because they did not show a preference for the novel vs familiar mouse (paired Student’s t-test, t=2.7, df=4, p=0.06). (I) The total exploration time during the social discrimination task was not significantly different between normal and epileptic mice (unpaired Student’s t-test, t=0.2, df=12, p=0.86).

Next, we used a two-choice social memory recognition task to determine whether chronic epileptic mice showed any impairment in social memory (Fig. 7E). Mice were allowed to explore two novel stimulus mice for 5 min in learning trial 1. Immediately thereafter, in learning trial 2, mice were allowed to explore the same stimulus mice for 5 min but in reversed position. Subsequently mice were returned to the home cage for 1 hr, and mostly slept during that time. After 1 hr, there was a recall test where one of the two stimulus mice (chosen at random) was presented as well as a novel mouse (Fig. 7E). Comparisons between discrimination scores suggested that normal mice (n=9) spent significantly more time exploring the novel mouse compared to the familiar mouse in the recall trial, indicative of social memory (One way ANOVA, F(5, 30)=32.17, p<0.0001; Fig. 7F). In contrast, the epileptic mice (n=5) were impaired in the task because they failed to show a significant discrimination of the novel and familiar mice (One way ANOVA, F(2, 12)=2.28, p=0.14; Fig. 7G). When we compared the times that mice spent exploring the familiar vs novel mouse in the recall test, we found significantly longer exploration of the novel mouse only for normal (Wilcoxon signed rank test, p=0.003; Fig. 7H), but not for epileptic mice (paired Student’s t-test, t=2.7, df=4, p=0.06; Fig. 7H). Importantly, the total exploration time during the task did not differ between the normal and the epileptic mice (unpaired Student’s t-test, t=0.2, df=12, p=0.86; Fig. 7I).

CA2 activity is required for SWRs,^12^ brief ~150-Hz oscillations observed during slow wave sleep that are needed for social memory consolidation.^13^ We therefore wondered whether the abnormal HFOs seen in the epileptic mice (e.g., Fig. 6) might have interfered with social memory consolidation. We did not suspect IIS or seizures were involved because they were relatively rare or absent before and during the task described above. In contrast, HFOs were common. To test the idea that HFOs interfered with social memory in epileptic mice, we recorded epileptic mice by video-EEG for 1 hr periods before the task (baseline-sleep), during exploration (Trial 1, Trial 2, recall test) and during the sleep period preceding the recall test (post-sleep; Fig. 8A). We found that the spectral power of HFOs was significantly different between the different phases of the task (One way ANOVA, F(2, 4)=8.38, p=0.03; Fig. 8B). Post-hoc comparisons confirmed that HFO power was significantly higher during the baseline-sleep vs exploration phases of the social discrimination task (p=0.04; Fig. 8B). Closed-loop silencing of HFOs during the rest period between learning Trial 2 and the recall test significantly reduced HFO power compared to the baseline-sleep period (p=0.04; Fig 8B). Remarkably, HFO disruption was sufficient to rescue the social memory deficit in the epileptic mice (One way ANOVA, F(2, 12)=6.01, p=0.01; Fig. 8C). Indeed, the mice spent significantly longer times exploring the novel vs familiar mouse during the recall test (paired Student’s t-test, t=4.3, df=4, p=0.04; Fig. 8D). When we compared discrimination scores between control, epileptic, and the closed-loop epileptic group during the recall test, we found a statistically significant main effect for group (One way ANOVA, F(2, 14)=37.35, p<0.0001; Fig. 8E). Post-hoc comparisons, confirmed that discrimination scores were not significantly different between the control and closed-loop group (p=0.99; Fig. 8E), suggesting restoration of social memory based on discrimination scores comparable to normal levels. Other post-hoc tests confirmed this result, showing the control group had significantly greater values than the epileptic group (p<0.0001; Fig. 8E), and the epileptic group was impaired compared to the closed-loop epileptic group (p<0.0001; Fig. 8E).

**Figure 8:**
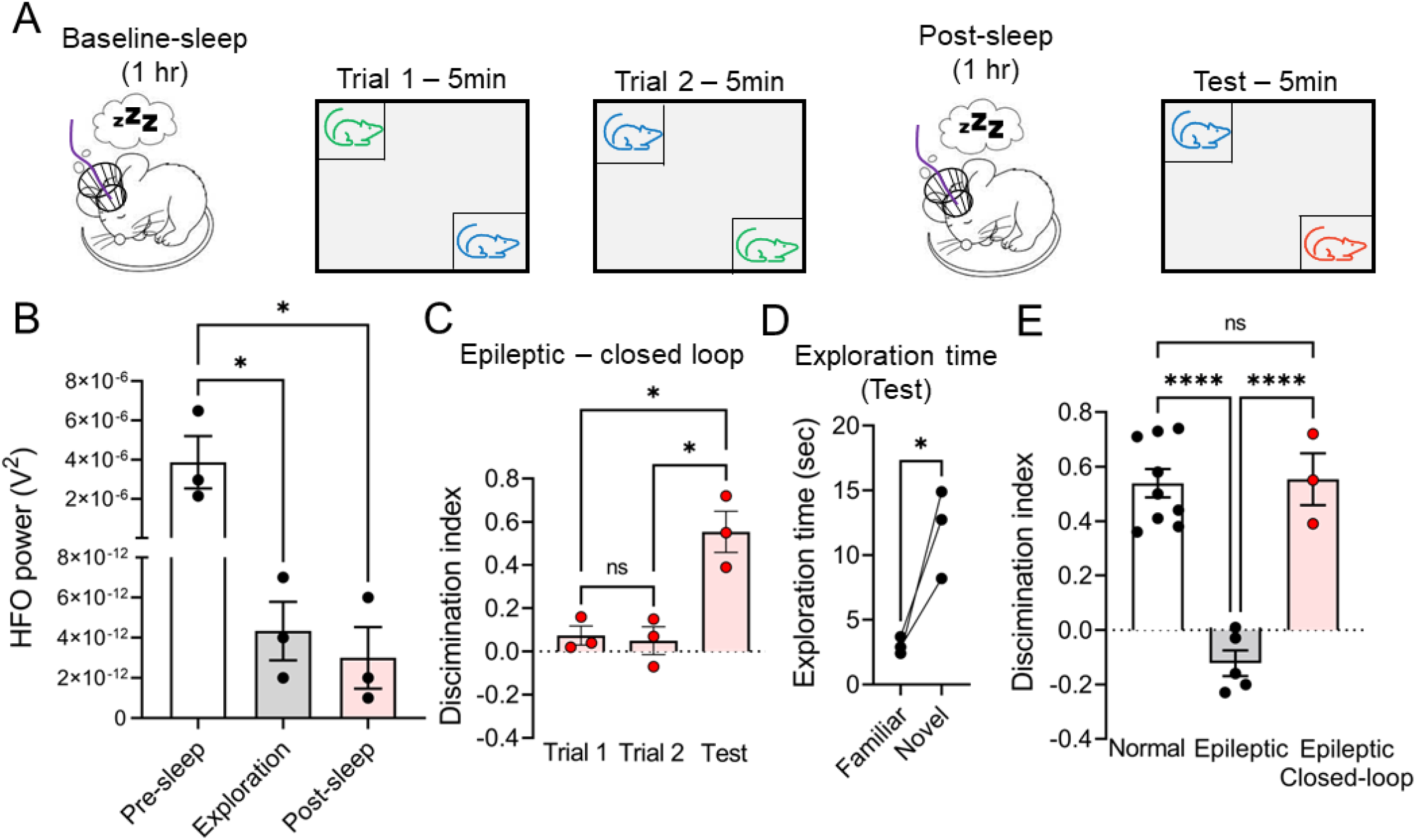
Closed-loop silencing of HFOs rescued social memory deficits of epileptic mice. (A) Diagram of the closed-loop social discrimination task. Mice were recorded by video-EEG continuously for 3 days preceding the task and during all phases of the task to ensure seizures did not occur which was the case for all mice tested. A baseline sleep period lasting for 1 hr preceded the task (baseline-sleep). After 1 hr, the subject mouse was presented with 2 mice placed on the opposite corners of an arena for 5 min during trial 1. During trial 2 mice are switched in position and the subject mouse was allowed to explore for 5 min. The subject mouse was placed back in its homecage for 1 hr during which closed-loop protocols were applied (post-sleep). After 1 hr the subject mouse was placed back in the arena where one mouse (familiar) was kept in its original position from trial 2 and a novel mouse was placed in the opposite corner of the arena. (B) HFO power was significantly different between baseline-sleep, exploration, and post-sleep periods (One way ANOVA, F(2, 4)=8.38, p=0.03). Post-hoc comparisons showed that HFO power was significantly reduced by closed-loop silencing of HFOs compared to the baseline-sleep period (p=0.04). There was higher HFO power during the baseline-sleep period compared to exploration (p=0.04), consistent with the prominence of HFOs in sleep. (C) Epileptic mice that underwent close-loop silencing of HFOs during the 1 hr interval in the homecage successfully recognized the novel mouse from the familiar mouse during the recall test (One way ANOVA, F(2, 4)=10.76, p=0.02; post-hoc test, novel vs familiar, p=0.01 during the recall test) (D) The time spent exploring the familiar vs novel mouse during the recall test trial was significantly longer (paired Student’s t-test, t=4.3, df=4, p=0.04). (E) Discrimination scores in the recall test. A statistically significant main effect for group was found (One way ANOVA, F(2, 14)=37.35, p<0.0001). Post-hoc comparisons confirmed that discrimination scores were significantly different between the control and epileptic group (p<0.0001), the epileptic and closed-loop group (p<0.0001), but not between the control and closed-loop group (p=0.99).

## DISCUSSION

### Summary of main findings

Our results both confirm and extend the findings that CA2 makes an important contribution to seizure activity in mouse models of TLE. Perhaps of even greater significance, we show for the first time that closed-loop manipulations of CA2 can rescue a social behavioral comorbidity of TLE. These main findings can be summarized as follows. First, we show that closed-loop acute CA2 silencing at the start of seizures significantly reduced hippocampal seizure duration and power. In addition, we found a robust reduction in cortical seizure power. Second, we show that the time spent in convulsive behavior is significantly reduced by CA2 silencing as seizures became less severe. Third, we found that CA2 silencing can abate IIS and HFOs in real-time. Last, we show that selective silencing of HFOs improves social recognition memory in a social discrimination task. These findings support for the first time an active role of CA2 PNs in several electrophysiological and behavioral hallmarks of TLE and also support the view that area CA2 could become a novel therapeutic target for TLE and its associated social comorbidities.

### CA2 PNs reduced hippocampal seizure duration and cortical seizures, as well as convulsions

Our findings show that CA2 activity directly contributes to seizure dynamics in TLE. Although CA2 PNs are known to become hyperexcitable in human TLE,^31^ and after PILO-SE^8^ and IHKA-SE^7^ in mice there has been only one study^8^ to date that inhibited CA2 selectively and showed spontaneous seizures were reduced in frequency. That study, however, used continuous chemogenetic inhibition of area CA2 and left open the question of whether specific targeting of CA2 at the onset of a seizure could abrogate it. Importantly, as prolonged CA2 silencing can have secondary effects on circuit activity^9^ that study could not distinguish a direct acute effect of CA2 silencing on seizure activity compared to an indirect effect. Moreover, the previous study did not address whether CA2 contributed to cortical seizure activity, IIS, HFOs, or any behavioral comorbidities.

The present study provides a significant advance as we now show that CA2 inhibition that starts at the onset of seizures truncates them, reduces cortical seizures, and reduces convulsive manifestations. These results suggest that CA2 is not just a passive regulator of seizure activity, but rather controls hippocampal seizures and their manifestations elsewhere. Furthermore, the results show that selective CA2 inhibition can truncate IIS and reduce HFOs. Remarkably, reducing HFOs in a social memory test led to the rescue of social memory impairment in epileptic mice. The results were obtained in two models of TLE and both males and females, supporting the robust nature of the findings.

### CA2 PNs can truncate interictal EEG abnormalities in TLE

Recordings from human brain slices suggested that the CA2 subfield can generate spontaneous activity in the form of IIS which is independent of other hippocampal areas.^30–32^ However, there has been no direct evidence showing that IIS can be controlled by CA2 *in vivo*. Here we show for the first time that CA2 can prevent IIS, which suggests that CA2 is necessary for IIS generation. This is important because CA2 inhibition may provide a new way to limit IIS from propagating to extra-hippocampal areas and disturbing function.

HFOs are considered another interictal EEG abnormality that can also impair memory.^33–35^ It has been hailed as an epilepsy biomarker, occurring in the context of various different epileptogenic etiologies such as human epilepsy,^36^ traumatic brain injury^37^, and in rodents, after IHKA-SE ^19^ and PILO-SE^16^. HFOs are best known as a biomarker of the epileptogenic zone and are often used in the presurgical evaluation of patients with TLE,^38^ although the topic has been debated.^39^

Our studies have focused on HFOs>250 Hz which are absent in normal human and rodent brain tissue and thus can be considered an abnormal electrical pattern.^16,19,40^ Despite their prominent role in epilepsy, no studies have attempted to selectively silence HFOs in real-time and at the location that they are recorded. Instead, prior studies have only focused on removing brain tissue where HFOs were presumably generated.^41^ Thus, our findings that HFOs can be selectively silenced only when and where they occur suggests for the first time that HFOs can be controlled, and CA2 is a critical contributor to HFOs.

### Selective silencing of CA2 HFOs improves social memory performance in epileptic mice

Hippocampal area CA2 is important for the synchronization of population activity during SWRs^12^ which are essential for the recall of a social experience.^13^ In TLE, when social memory is impaired, HFOs>250 Hz also may be critical because they are implicated in cognitive dysfunction of humans with TLE^33^ as well as rodent models.^34,35^ However, no studies have tested social memory and none have asked whether selective targeting of HFOs could rescue social memory. Here we show for the first time that selective reduction of HFOs during the interval between learning and recall of a social experience is sufficient to improve social memory in chronic epileptic mice. The phase between learning and recall is significant because it is when memory consolidation in response to SWRs occurs. Although CA2 silencing can disrupt social memory in the normal rodent,^13^ epileptic mice show hyperexcitability in CA2,^8^ so silencing of HFOs could contribute to the reduction of this hyperexcitability and therefore return CA2 excitability closer to normal levels.

How CA2 regulates social memory in epilepsy is not clear at the present time. Our data support a defect in the consolidation of a social experience. However, our experiments cannot exclude an effect on encoding. In epilepsy, there could also be an inappropriate communication between CA2 and its connections, such as CA1^11^ which could occur during encoding, consolidation, retrieval, or all phases of social learning and memory. Other possibilities include regulation by CA2 interneurons which are disrupted in the PILO model of TLE.^42^ CA2 interneurons are a logical candidate to regulate social memory in TLE because of the evidence they are involved in social memory deficits in schizophrenia.^43^

### Could hippocampal area CA2 become a novel therapeutic target for human TLE?

Previous studies looking at the role of different hippocampal areas in reducing seizures in TLE mainly focused on relatively vulnerable areas in TLE such as the DG hilus,^44^ which undergoes extensive neurodegeneration in the majority of TLE cases.^5^ Thus, it may be challenging to target this area in the human epileptic brain. As an alternative, other studies have suggested that extra-hippocampal areas such as the thalamus,^45,46^ the medial septum,^47^ the cerebellum^48^ could also be targeted. Therefore, targeting areas that are resistant in the hippocampus are also valuable, such as area CA2.

Hippocampal area CA2 could become a novel TLE target for several reasons. First, CA2 PNs are characterized by a unique molecular profile^49^ and pattern of expression of G protein-coupled receptors, intracellular signal transduction proteins, and ion channels, which could potentially allow CA2 to be targeted selectively through pharmacological agents. Second, we have shown that CA2 PNs not only control seizures but also social memory impairments. Thus, their selective targeting may provide a dual benefit. An intriguing possibility is to use CA2 stimulation in patients with TLE. Although stimulation at one time might excite CA2 neurons, patterned stimulation to inhibit brain areas can be therapeutic.^50^ Such therapeutic options offer promising new avenues to treat the approximately 30% of TLE patients that are resistant to current treatments, as well as the majority of patients that do respond to medication but suffer serious side effects.

## CONCLUSIONS

This study showed that selective closed-loop CA2 silencing in two mouse models of TLE reduces hippocampal seizure duration, cortical seizure power, and convulsions, as well as IIS and HFOs. Furthermore, inhibiting CA2 HFOs during the interval between learning and recall of a social experience rescued the impairment in this test in seizure-untreated epileptic mice. The results provide a significant advance in our understanding of area CA2 and its powerful role in epilepsy and its social comorbidities.

## Abbreviations

DG: Dentate gyrus
EEG: Electroencephalography
HFOs: High frequency oscillations
IIS: Interictal spikes
PNs: Pyramidal neurons
TLE: Temporal lobe epilepsy

## ACKNOWLEDGMENTS

We thank the members of the Scharfman and Siegelbaum labs for fruitful discussions. This work was supported by National Institutes of Health grant R01 NS-106983 from NIH (PIs, H.E.S. and S.A.S.).

